# NhaR, LeuO and H-NS are part of an expanded regulatory network for ectoine biosynthesis expression

**DOI:** 10.1101/2022.11.11.516238

**Authors:** Katherine E. Boas Lichty, Gwendolyn J. Gregory, E. Fidelma Boyd

## Abstract

Bacteria accumulate compatible solutes, to maintain cellular turgor pressure when exposed to high salinity. In the marine halophile *Vibrio parahaemolyticus,* the compatible solute ectoine is biosynthesized *de novo*, which is energetically more costly than uptake; therefore, tight regulation is required. To uncover novel regulators of the ectoine biosynthesis *ectABC-asp_ect* operon, a DNA affinity pulldown of proteins interacting with the *ectABC-asp_ect* regulatory region was performed. Mass spectrometry analysis identified, amongst others, three regulators: LeuO, NhaR, and the nucleoid associated protein H-NS. In-frame non-polar deletions were made for each gene and P*_ectA_-gfp* promoter reporter assays were performed in exponential and stationary phase cells. P*_ectA_-gfp* expression was significantly repressed in the Δ*leuO* mutant and significantly induced in the Δ*nhaR* mutant compared to wild type, suggesting positive and negative regulation, respectively. In the Δ*hns* mutant, P*_ectA_-gfp* showed increased expression in exponential phase cells, but no change compared to wild type in stationary phase cells. To examine whether H-NS interacts with LeuO or NhaR at the ectoine regulatory region, double deletion mutants were created. In a Δ*leuO/*Δ*hns* mutant, P*_ectA_-gfp* showed reduced expression, but significantly more than Δ*leuO* suggesting H-NS and LeuO interact to regulate ectoine expression. Whereas Δ*nhaR/*Δ*hns* had no additional effect as compared to Δ*nhaR* suggesting NhaR regulation is independent of H-NS. To examine *leuO* regulation further, a P*_leuO_-gfp* reporter analysis was examined that showed significantly increased expression in the Δ*leuO*, Δ*hns* and Δ*leuO/*Δ*hns* mutants as compared to wild type, indicating both are repressors. Growth pattern analysis of the mutants in M9G 6%NaCl, showed growth defects compared to wild type, indicating that these regulators play an important physiological role in salinity stress tolerance.

**Importance:** Ectoine is a commercially used compatible solute that acts as a biomolecule stabilizer because of its additional role as a chemical chaperone. A better understanding of how the ectoine biosynthetic pathway is regulated in natural bacterial producers can be used to increase efficient industrial production. The *de novo* biosynthesis of ectoine is essential for bacteria to survive osmotic stress when exogenous compatible solutes are absent. This study identified LeuO as a positive regulator and NhaR as a negative regulator of ectoine biosynthesis and also showed that similar to enteric species, LeuO is an anti-silencer of H-NS. In addition, defects in growth in high salinity among all the mutants suggest that these regulators play a broader role in the osmotic stress response beyond ectoine biosynthesis regulation.

## Introduction

When bacteria are exposed to high salinity conditions, an efflux of water across the osmotic gradient occurs lowering cellular turgor pressure (1–5). The initial short-term response to osmotic up shock is the uptake of potassium (K+), which is accompanied by glutamate accumulation in Gram-negative bacteria to maintain electro neutrality. This is the short-term strategy because high concentrations of K+ have deleterious effects on cellular processes (1, 2, 6, 7). The long term response to increased osmotic stress is characterized by the uptake or biosynthesis of compatible solutes, which are small organic molecules that provide osmotic protection by helping to maintain the turgor pressure of the cell and can be accumulated to extremely high intracellular concentrations (3, 8, 9).

The compatible solute ectoine was initially discovered to provide osmotic protection in anoxygenic phototrophs (10), but subsequent analysis showed that it functions as a compatible solute in numerous Gram-negative and Gram-positive bacteria (reviewed in (3, 8, 11–14), which included many members of the family *Vibrionaceae* (15–19). Accumulation to very high intracellular concentrations without interfering with molecular processes is a hallmark of compatible solutes. Additionally, ectoine acts as a chemical chaperone through the stabilization of proteins, thus it is an important compound for commercial uses in medicine and cosmetics to stabilize biomolecules against factors such as heating, freezing, desiccation, and UV radiation (20–23). Therefore, understanding the regulation of the ectoine biosynthetic pathway is important to increase efficient industrial production using natural bacterial producers (24).

Biosynthesis of ectoine (1,4,5,6-tetrahydro-2-methyl-4-pyrimidinecarboxylic acid) is *de novo* from L-aspartic acid, and performed by the evolutionarily conserved operon *ectABC,* which encodes the EctA, EctB, and EctC proteins (25–27) (*Fig. S1*). In many species an aspartokinase (*asp_ect*) is clustered with the ectoine biosynthesis operon, which converts aspartic acid to β-aspartyl phosphate, which is then converted to L-aspartate-β-semialdehyde by aspartate semialdehyde dehydrogenase (Asd). This intermediate from the aspartic acid pathway is incorporated into the ectoine biosynthesis pathway when it is converted to L-2,4-diaminobutyrate by EctB (L-2,4-diaminobutyrate transaminase). EctA (L-2,4-diaminobutyrate N^γ^-acetyltransferase) then acetylates this product to form N^γ^-acetyldiaminobutyrate. The final step to produce ectoine is a cyclic condensation reaction performed by EctC (L-ectoine synthase) (13, 27).

In some species, clustered with the ectoine biosynthesis operon is *ectR*, a MarR-type regulator shown to repress transcription of the ectoine operon (24, 28–30). Among *Vibrionaceae*, an *ectR* homolog is present only in *Aliivibrio* species (31, 32). Previous studies in *V. parahaemolyticus* and *V. cholerae* showed that *ectABC-asp_ect* transcription was repressed in low salinity by another MarR-type regulator named CosR (17, 31, 33). CosR from *V. parahaemolyticus* showed 31% amino acid identity to *Aliivibrio fischeri* EctR, with less than 60% query coverage and does not cluster with *ectABC-asp_ect*, but is located elsewhere in the genome (31). The quorum sensing master regulators AphA and OpaR activate and repress expression of *ectABC-asp_ect*, respectively in *V. parahaemolyticus* (17). AphA and OpaR also regulate expression of CosR, creating a tightly controlled feed-forward loop for ectoine expression across the growth curve (17).

In this study, we set out to identify novel regulators of the ectoine biosynthesis genes by performing a DNA affinity chromatography pulldown using the regulatory region of the *ectABC-asp_ect* operon. We identified numerous candidates and selected the regulators LeuO, NhaR, OmpR, TorR, and the nucleoid associated protein (NAP) H-NS to examine further. We constructed in-frame nonpolar deletions in each gene. These regulators play a role in the regulation of the ectoine biosynthesis genes based upon P*_ectA_-gfp* expression reporter assays. In addition, analysis of double mutants Δ*leuO/Δhns* and Δ*nhaR/Δhns* showed that H-NS interacted with LeuO but not NhaR to control ectoine gene expression. Importantly, when these deletion mutants were grown under high salinity conditions a growth defect was observed, suggesting that the regulators NhaR, LeuO, and H-NS are physiologically important for growth in high salinity conditions.

## Results

### Ectoine is biosynthesized in cells grown in M9G 3%NaCl, but absent in M9G 1%NaCl

To confirm *V. parahaemolyticus de novo* ectoine biosynthesis is a response to increased salinity, ^1^H nuclear magnetic resonance spectroscopy (^1^H-NMR) at 600MHz in the presence of deuterium oxide as the solvent was performed. Ethanol extracts of *V. parahaemolyticus* wild type cells grown overnight at 37°C in minimal medium supplemented with glucose (M9G) with either 1%NaCl or 3%NaCl were used to obtain ^1^H-NMR spectra (*Fig. S2A*). Peaks corresponding to the hydrogen atoms of ectoine are labeled and were present in M9G 3%NaCl extracts but were not detectable in M9G 1%NaCl extracts as previously shown (15) (*Fig. S2A*). In addition to intracellular accumulation of ectoine, we also wanted to demonstrate that *ectABC-asp_ect* expression is increased in M9G 3%NaCl compared to 1% NaCl. A transcriptional reporter assay was performed, where *gfp* was placed under the control of the ectoine regulatory region (P*_ectA_-gfp*). The expression was examined in wild type cells grown in either M9G 1%NaCl or M9G 3%NaCl by measuring the specific fluorescence of P*_ectA_-gfp* as a cumulative readout of *ectA* transcription (*Fig. S2B*). The observed level of specific fluorescence of P*_ectA_-gfp* was significantly increased in M9G 3%NaCl (4.0-fold) relative to that of cells grown in M9G 1%NaCl (*Fig. S2B*).

### Identification of putative regulators of the ectoine biosynthesis operon

With the knowledge that ectoine is present in cells grown in M9G 3%NaCl, we performed a promoter pulldown under this growth condition. A promoter pulldown was performed using a 5’-biotin tagged 323-bp DNA fragment of the *ectABC-asp_ect* regulatory region bound to streptavidin beads. As a negative control DNA probe, the coding region of the *ectB* gene was used. To identify novel regulators of ectoine biosynthesis, we grew cells to mid-exponential phase in M9G 3%NaCl and subsequent cell lysate was incubated with DNA probe-coated beads. Bound proteins were then eluted using a stepwise NaCl gradient and separated with SDS-PAGE gel (*Fig. S3A*). Bands that were present in the target lanes, but not present in the *ectB* negative control lanes were selected for analysis via mass spectrometry (*Fig. S3B*). The proteins identified with mass spectrometry analysis were aligned to the *V. parahaemolyticus* genome and sorted by gene class and ranked by score (*Table S1*). The mass spectrometry score is the combined sum of all scores for an observed spectra that can be matched to an individual peptide. The higher the score the more confident the match to the specified peptide. Candidate proteins of interest that belonged to the transcriptional regulator gene class were selected. These candidate proteins are described in Table S1 and include homologs of regulators NhaR (VP0527), OmpR (VP0154), and TorR (VP1032). NhaR and OmpR were shown in *Escherichia coli,* which does not produce ectoine, to be involved in the osmotic stress response (34–39). *V. parahaemolyticus* NhaR shows 62%, OmpR 84%, and TorR 57% amino acid identities to homologs in *E. coli*. NhaR, OmpR, and TorR have not been examined in *Vibrio* species. An additional regulators of interest, LeuO (VP0350), a LysR-type regulator and H-NS (histone-like nucleoid structuring) protein were also examined further. In *V. parahaemolyticus,* LeuO was previously shown to be positively regulated by ToxR and is essential for acid stress tolerance (40, 41). H-NS in *E. coli* and *Salmonella enterica,* is a universal silencer of gene expression (42–47). In addition, an *E. coli* study by Shimada and colleagues identified 140 LeuO-binding sites with over 90% of these sites also showing H-NS binding. Further studies have shown that LeuO plays a significant role as an antagonist of H-NS gene silencing (42, 48, 49). While not chosen for examination in this study, two additional regulators did stand out for possible future analysis, VPA0036 (accession number Q87K62) and VP0489 (accession number Q87SD5) (*Table S1*). VPA0036 gave a high score, but is annotated as a hypothetical protein with no conserved domains. HHPred analysis uncovered a putative N-terminal winged-helix protein domain with a probability of 96.53% and E-value 0.016. VP0489 is annotated as an aerobic respiration control protein FecA and ArcA a two-component system response regulator.

As a proof of concept for the pulldown, LeuO and NhaR proteins were purified and visualized on SDS-PAGE gels showing an expected 36.2 kDa LeuO protein and a 33.9 kDa NhaR protein (*Fig. S4*). Electrophoretic mobility shift assays (EMSAs) were performed first using the 323-bp regulatory region of *ectABC-asp_ect* as a DNA probe (*Fig. S5A*). For the LeuO EMSA, we used the regulatory region of the *leuO* gene as a positive control and show binding in a concentration dependent manner for this and also for the P*ectA* probe 1 (*Fig. S5B*). Binding of NhaR to P*ectA* was also confirmed in the EMSA (*Fig. S5C*). To examine binding of LeuO and NhaR further, probe 1 was divided into three similarly sized probes P*ectA* 1A, 1B and 1C. Purified protein LeuO showed band shifts for probe 1A and a complete shift for probe 1C (*Fig. S5D*). For NhaR all three probes showed shifts in a concentration dependent manner (*Fig. S5E*).

### PectA-gfp reporter assays shows differential expression in ΔleuO, ΔnhaR, ΔompR and Δ*torR mutants*

To examine the role of *leuO*, *nhaR*, *ompR*, and *torR* candidate regulators, we performed GFP reporter assays using a reporter plasmid where *gfp* was under the control of the ectoine regulatory region (P*_ectA_-gfp*). This reporter plasmid was transformed into wild type or in-frame non-polar mutants of Δ*leuO*, Δ*nhaR*, Δ*ompR*, and Δ*torR*. Cultures were grown overnight in M9G 3% NaCl and relative fluorescence intensity (RFU) was measured. Specific fluorescence was calculated by dividing RFU by OD. From this analysis, the relative fluorescence (RFU) measured in the Δ*leuO* mutant background was 1.7-fold reduced as compared to wild type (*Fig. 1*). This data suggests that LeuO is an activator of *ectABC-asp_ect*. The relative fluorescence measured in a Δ*nhaR* background was significantly increased 1.8-fold as compared to wild type, suggesting that NhaR is a repressor of *ectABC-asp_ect* (*Fig. 1*). P*_ectA_-gfp* expression in the Δ*ompR* (1.3-fold) and Δ*torR* (1.4-fold) mutants were also increased as compared to wild type, but not to the same level as NhaR.

**Fig. 1.**
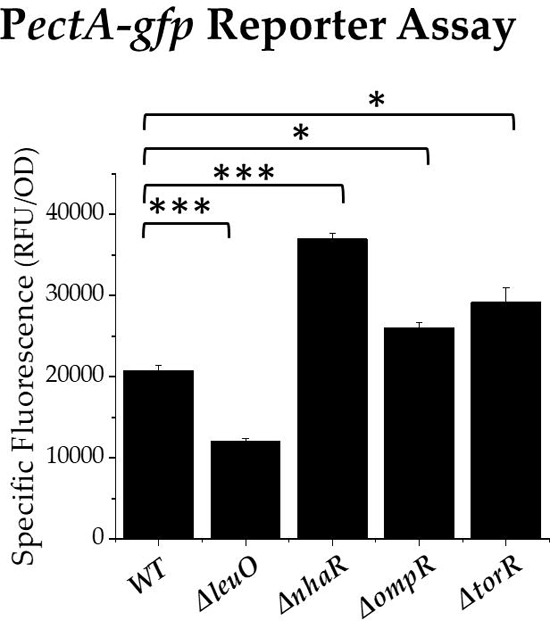
Expression of a P*_ectA_-gfp* transcriptional fusion in wild-type (WT) *V. parahaemolyticus* Δ*leuO,* Δ*nhaR*, Δ*ompR,* and Δ*torR* mutants. Cultures were grown overnight in M9G 3% NaCl and relative fluorescence intensity (RFU) was measured. Specific fluorescence was calculated by dividing RFU by OD. Mean and standard deviation of three biological replicates are shown. Statistics were calculated using a Student’s t-test; (*, P < 0.05; ***, P < 0.001).

### LeuO and H-NS interplay to control PectA-gfp

In *E. coli* and *V. cholerae*, H-NS has been shown to interact with LeuO in both cooperative and antagonistic relationships at different gene loci (42, 50–53). Therefore, we wanted to examine whether H-NS and LeuO interacted at the *ectABC-asp_ect* regulatory region. We constructed an in-frame non-polar Δ*hns* mutant and a double Δ*leuO/Δhns* mutant. In exponential phase cells (OD ∼ 0.4), P*_ectA_-gfp* in the Δ*leuO* mutant showed significant reduced expression, and in the Δ*hns* mutant expression was significantly increased (*Fig. 2A*). However, in the Δ*leuO/*Δ*hns* double mutant P*_ectA_-gfp* expression was reduced, but not to the same level as the Δ*leuO* single mutant (*Fig. 2A*). In stationary phase cells (OD ∼ 1.0), P*_ectA_-gfp* relative fluorescence was similar to exponential cells in all strains except the Δ*hns* mutant, which was similar to wild-type (*Fig. 2B*). These data indicate interplay between both LeuO and H-NS to control ectoine gene expression with LeuO playing an anti-silencer role.

**Fig. 2.**
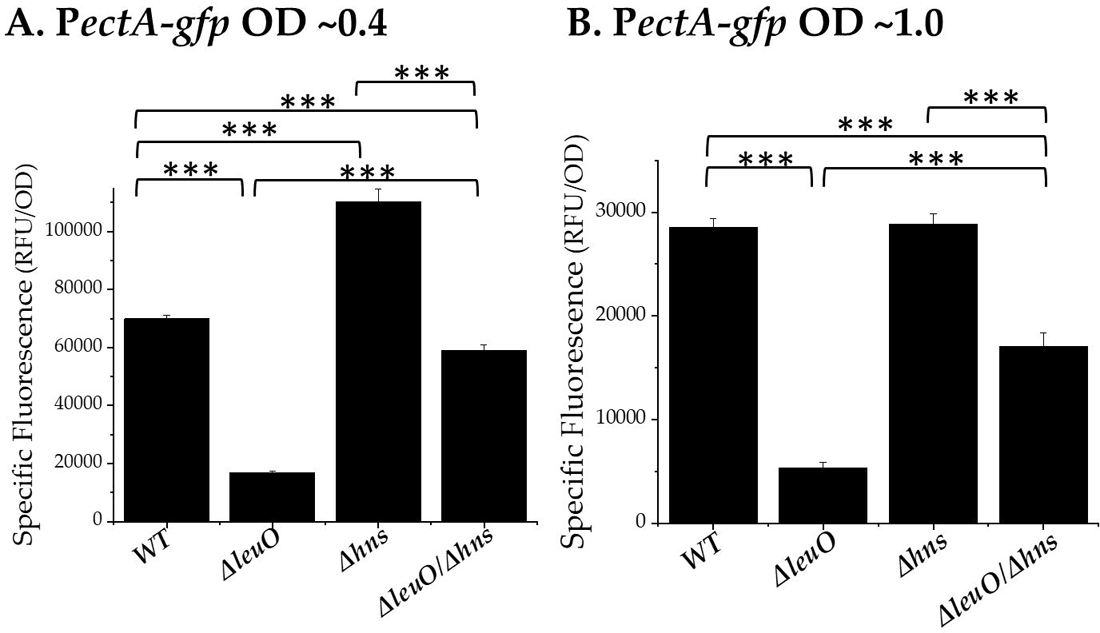
Expression of a P*_ectA_-gfp* transcriptional fusion in wild-type (WT) *V. parahaemolyticus* Δ*leuO*, Δ*hns,* and Δ*leuO/*Δ*hns* mutants. Cultures were grown *A*. to OD ∼0.4 or *B*. OD ∼1.0 in M9G 3%NaCl and relative fluorescence intensity (RFU) was measured. Specific fluorescence was calculated by dividing RFU by OD. Mean and standard deviation of three biological replicates are shown. Statistics were calculated using an ANOVA followed by Tukey-Kramer post hoc test; (***, P < 0.001).

To further investigate the relationship between LeuO and H-NS in *V. parahaemolyticus*, a *gfp*-expressing reporter plasmid under the control of the LeuO regulatory region (P*_leuO_-gfp*) was constructed. Relative fluorescence was measured in cells grown to OD ∼1.0 in M9G 3%NaCl in wild type, Δ*leuO,* Δ*hns,* Δ*leuO/*Δ*hns,* and Δ*toxR*. When compared to wild type, P*_leuO_-gfp* expression in the Δ*hns* mutant showed a 4.0-fold increased relative fluorescence indicating that H-NS is a repressor of *leuO* expression (*Fig. 3*). P*_leuO_-gfp* expression was 2.4-fold higher in Δ*leuO* and 2.8-fold higher in Δ*leuO/*Δ*hns* when compared to wild type. In *V. parahaemolyticus,* ToxR is a positive regulator of *leuO* expression (40, 54, 55) and this was confirmed in this reporter assay, with Δ*toxR* showing reduced P*_leuO_-gfp* (1.8-fold) expression when compared to wild type (*Fig. 3*). These data indicate that both H-NS and LeuO are repressors of *leuO* expression and in the absence of both negative regulators other proteins such as ToxR are likely able to access the regulatory region.

**Fig. 3.**
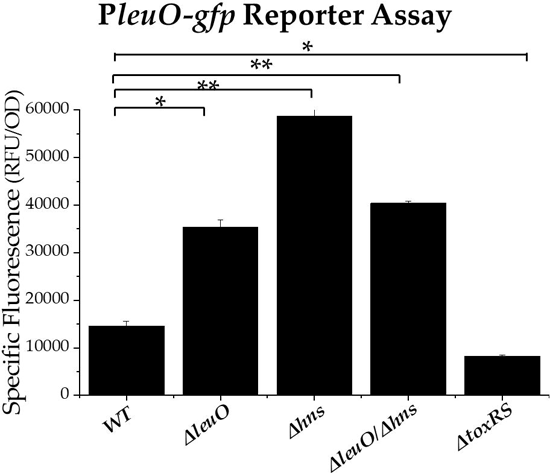
Expression reporter assays of P*_leuO_-gfp* transcriptional fusion in *V. parahaemolyticus* wild type (WT), Δ*leuO*, Δ*hns,* Δ*leuO/*Δ*hns, and* Δ*toxRS* mutants. Cultures were grown to OD ∼1.0 in M9G 3%NaCL and relative fluorescence intensity (RFU) was measured. Specific fluorescence was calculated by dividing RFU by OD. Mean and standard deviation of two biological replicates are shown. Statistics were calculated using a Student’s t-test; (*, P < 0.05; **, P < 0.005).

### H-NS and LeuO are important for growth in high salinity

To assess whether the deletions of *leuO* and *hns* have a physiological effect, we examined the ability of *V. parahaemolyticus* wild type, Δ*leuO,* Δ*hns,* and Δ*leuO/*Δ*hns* mutants to grow in M9G 1% NaCl, 3%NaCl, or 6%NaCl at 37°C. In M9G 1%NaCl and 3%NaCl, there were no significant growth defects observed among most strains, with the exception of Δ*leuO,* which reached a lower final OD (0.44) as compared to wild type (0.57) (*Fig. S6A, 6B*). Growth in M9G 6%NaCl showed the wild type and Δ*hns* had a 5 h lag phase, but the Δ*hns* mutant reached a lower final OD (0.37) as compared to wild type (0.54) (*Fig. 4).* The Δ*leuO* mutant had a lag phase of 6 h and Δ*leuO/*Δ*hns* had a 13 h lag phase. In addition, the double deletion mutant had a final OD595 of 0.2, which was significantly less than wild type with final OD of 0.54 (*Fig. 4*). Complementation of Δ*leuO* with pBA*leuO* and Δ*hns* with pBA*hns,* restored growth to wild type levels (*Fig. 4*). However, we were unable to complement the double mutant with either pBA*leuO* or pBA*hns*. Overall, these data demonstrate that both LeuO and H-NS play a significant role in the osmotic stress response and are required for optimal growth in high salinity.

**Fig. 4.**
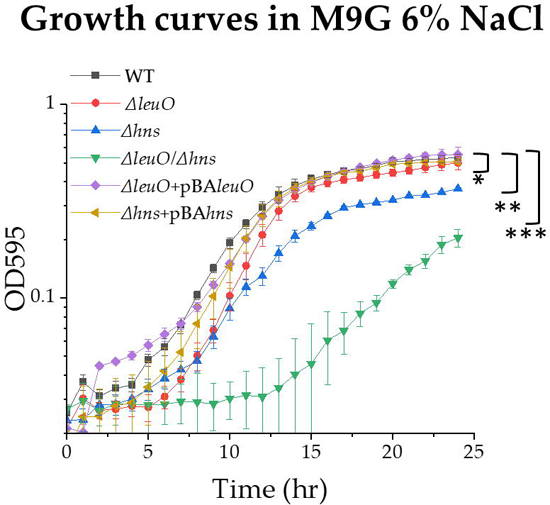
Growth analysis of Δ*leuO, Δhns* and *ΔleuO/Δhns* mutants in M9G 6%NaCl. Cells were grown to OD 0.5 in M9G 1%NaCl and then inoculated into M9G 6%NaCl and growth was measured every hour for 24h at 37°C. Mean and standard deviation of two biological replicates are shown. Also shown, complementation of Δ*leuO* with pBA*leuO* and Δ*hns* with pBA*hns*. For all growth curves the area under the curve (AUC) was calculated for each strain and a Student’s t-test was performed (*, P < 0.05; **, P < 0.005; ***, P < 0.001).

### NhaR is a negative regulator of the ectoine biosynthesis operon

Next, we considered whether there was interplay between NhaR and H-NS in the regulation of *ectABC-ask*. To accomplish this, we examined P*_ectA_-gfp* in Δ*nhaR*, Δ*hns* and a Δ*nhaR/*Δ*hns* double mutant in M9G 3%NaCl in exponential and stationary phase cells (*Fig. 5*). In exponential cells (∼OD 0.4). P*_ectA_-gfp* expression was higher in Δ*nhaR* (1.8-fold) as compared to wild type confirming NhaR as a negative regulator (*Fig. 5A*). P*_ectA_-gfp*expression in the Δ*hns* and Δ*hns/*Δ*nhaR* mutants were both similarly increased compared to wild type (*Fig. 5A*). Examination of stationary phase cells (OD ∼ 1.0) gave similar results to expression in exponential cells with the exception of Δ*hns*, which showed P*_ectA_-gfp* expression levels similar to wild type (*Fig. 5B*). Overall, the data suggest that there is no interplay between H-NS and NhaR in ectoine expression because the double mutant was not significantly different than the single mutants. Next, we examined the effect of deleting *nhaR, hns,* and *hns/nhaR* on growth of *V. parahaemolyticus* in high salinity. Growth of the deletion mutants in M9G 1%NaCl and 3% NaCl was similar to wild type indicating no overall defect (*Fig. S6C, D*). However, Δ*nhaR,* Δ*hns,* and Δ*hns/*Δ*nhaR* mutants grown in M9G 6%NaCl, reached a final OD595 of 0.37, 0.36, and 0.38 respectively as compared to wild type OD595 of 0.5 (*Fig. 6*). Complementation of Δ*nhaR* with pBA*nhaR* restored growth to wild type levels (*Fig. 6*). These data indicates that there is a growth defect in high salinity for Δ*nhaR,* but deletion of *hns* does not contribute further to this defect. This supports the expression data that NhaR and H-NS do not interact to control ectoine expression. The data does indicate that NhaR is important for high salinity tolerance.

**Fig. 5.**
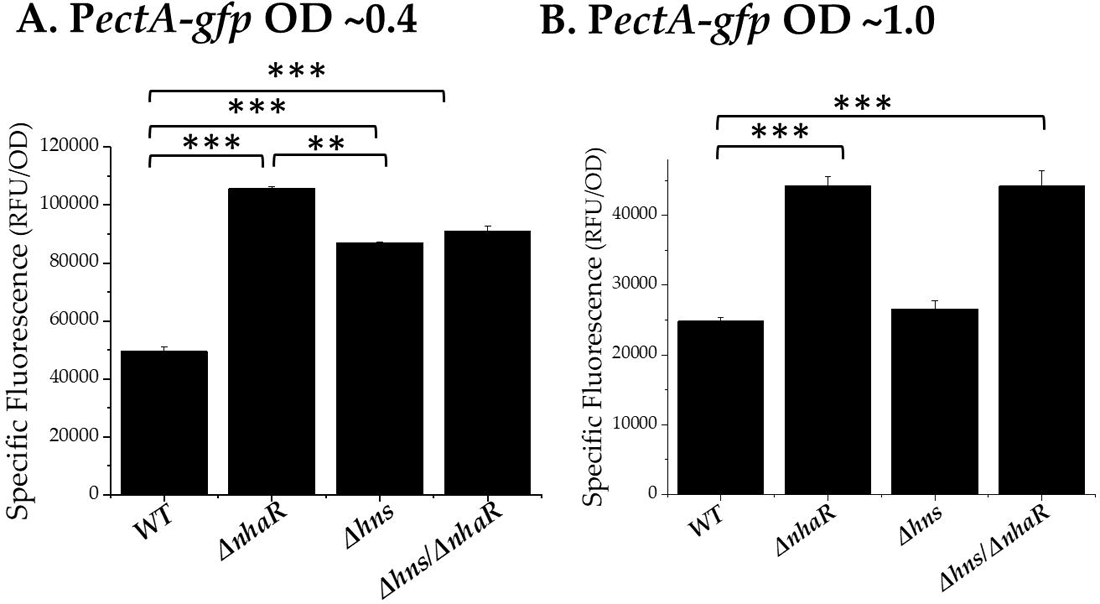
Expression of a P*_ectA_-gfp* transcriptional fusion in wild type (WT) *V. parahaemolyticus,* Δ*nhaR,* Δ*hns, and* Δ*hns/*Δ*nhaR* mutants. Cultures were grown *A*. to OD ∼0.4 or *B*. OD ∼1.0 in M9G 3%NaCl and relative fluorescence intensity (RFU) was measured. Specific fluorescence was calculated by dividing RFU by OD. Mean and standard deviation of three biological replicates are shown. Statistics were calculated using an ANOVA followed by Tukey-Kramer post hoc test; (***, P < 0.001).

**Fig. 6.**
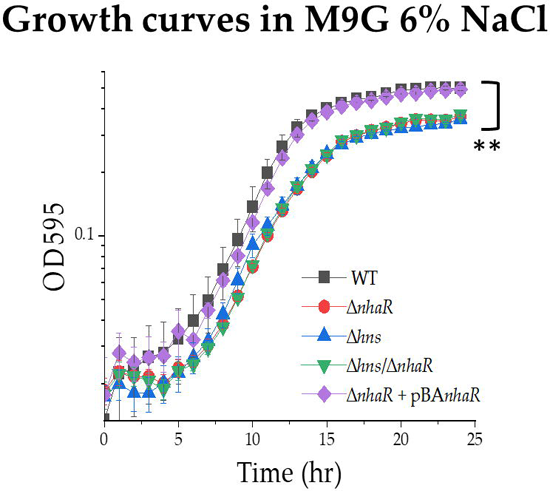
Growth analysis of Δ*nhaR, Δhns* and *ΔnhaR/Δhns* mutants in M9G 6%NaCl. Cells were grown to OD 0.5 in M9G 1%NaCl and then inoculated into M9G 6%NaCl and growth was measured every hour for 24 h at 37°C. Mean and standard deviation of two biological replicates are shown. Also shown, complementation of Δ*nhaR* with pBA*nhaR*. For all growth curves the area under the curve (AUC) was calculated for each strain and a Student’s t-test was performed (**, P < 0.005).

## Discussion

The accumulation of compatible solutes occurs either via uptake or biosynthesis and in *V. parahaemolyticus* there are six transporters and two biosynthesis operons for compatible solutes present (19, 56). The biosynthesis of the compatible solute ectoine and not glycine betaine was previously shown to be essential for *V. parahaemolyticus* survival in high salinity (15). However, ectoine biosynthesis is costly as it drains intracellular pools of metabolites such as aspartic acid and multiple regulators that provide tight control are needed (57).

In this study, we identified several new regulators of ectoine biosynthesis (*Fig. 7*). We found that NhaR acted as a repressor of ectoine biosynthesis. However, deletion of this repressor, showed a defect when grown under high salinity conditions (6% NaCl). This could be explained by the fact that NhaR is known to regulate genes related to the osmotic stress response outside of ectoine biosynthesis (58–61). In *E. coli*, *nhaR* is present in an operon with *nhaA,* which encodes a Na+/H+ antiporter. NhaR was shown to activate transcription of *nhaA*, which is induced in high salt and alkaline pH and is essential for survival under these conditions. It was also suggested that NhaR is a sensor and transducer of Na+ because the presence of Na+ alters the contact points between NhaR and the *nhaA* regulatory region (62). A homolog of *nhaA* is present in *V. parahaemolyticus*, and thus may also be activated by NhaR, which could explain the growth defect in high salinity for the *V. parahaemolyticus ΔnhaR* mutant.

**Fig. 7.**
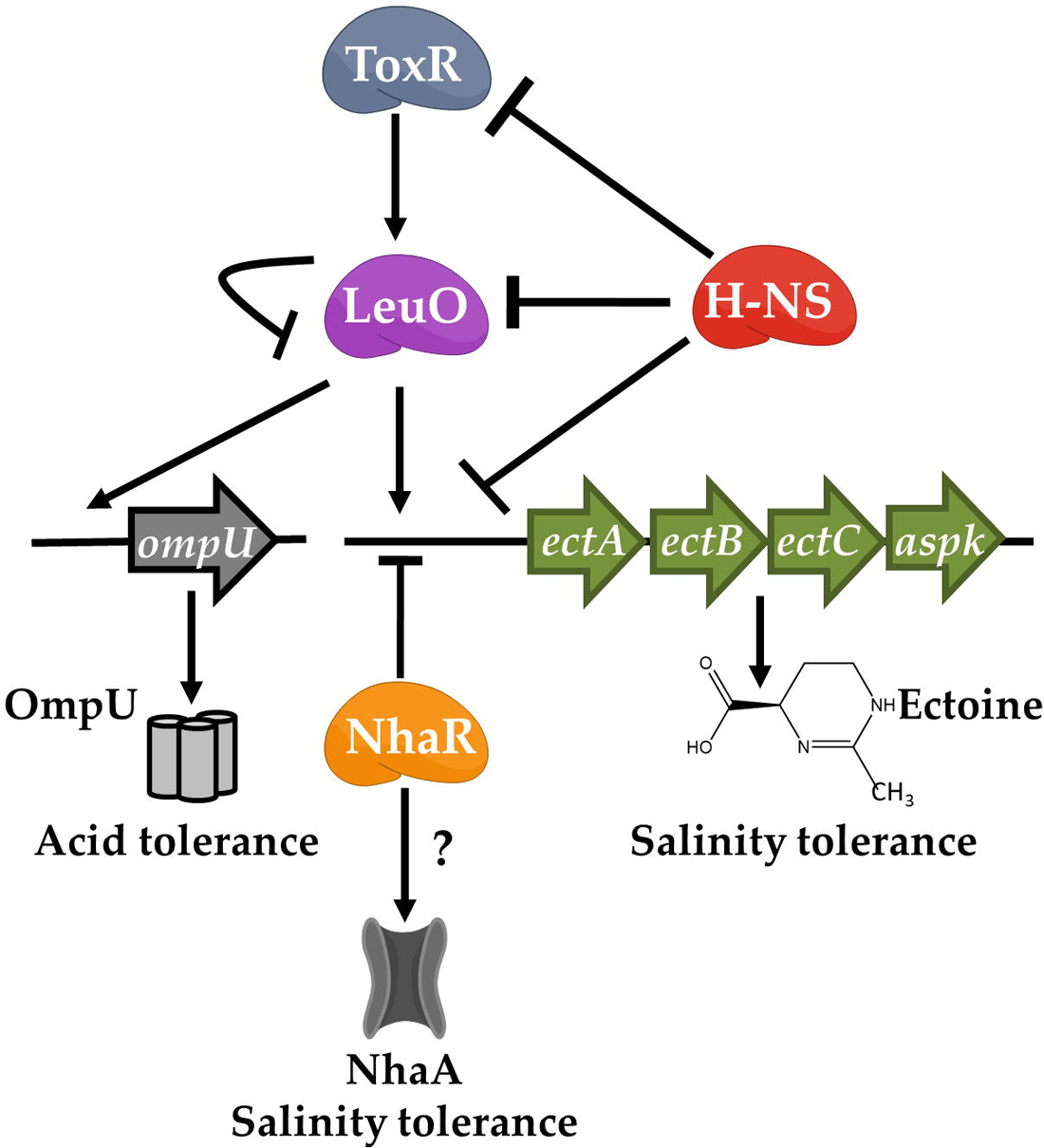
Model of ectoine biosynthesis gene expression in *V. parahaemolyticus*. Solid arrows represent positive regulation while hammer lines represent negative regulation. Previously it was shown that LeuO is required for OmpU expression, an important component of the acid stress response. H-NS was also shown to negatively regulated ToxR. In *E. coli*, NhaR is an activator of NhaA a symporter required for NaCl tolerance. Theindicates that this has not been shown in *V. parahaemolyticus*. Our data show that the NhaR and LeuO regulate *ectABC-asp_ect* and that H-NS is also part of this regulatory circuit.

Our data also established that LeuO is a positive regulator of ectoine biosynthesis gene expression and H-NS acts as a gene silencer. The analysis of a double Δ*leuO/*Δ*hns* mutant suggested that LeuO also likely plays a role as an anti-silencer of H-NS at this locus. In addition, we found that LeuO and H-NS both act as repressors of *leuO* in *V. parahaemolyticus*, further supporting the interaction between these two regulators. LeuO was first identified as a LysR-type regulator and shown to positively regulate the *leuABCD* leucine synthesis operon in *Salmonella enterica* (63, 64). Subsequently, chromatin-immunoprecipitation-on-chip analysis showed 178 LeuO binding sites in *S. enterica*, indicating its role as a global regulator (65). H-NS was also shown to be a global regulator in this species and in *E. coli*, silencing the transcription of a large number of horizontally acquired genes (46, 47, 66). In *V. parahaemolyticus*, LeuO was first named CalR as it was negatively regulated by calcium (41). LeuO was shown to positively regulate *ompU*, which encodes an outer membrane porin important for acid stress (*Fig. 7*) (40). Additionally, the global regulator ToxR was shown to be a positive regulator of *leuO* expression and deletion of either *leuO* or *toxR* resulted in a growth defect in acid conditions (40). In our study, *leuO* showed significantly increased expression in the *ΔleuO* mutant whereas in the *ΔtoxR* mutant *leuO* expression was reduced showing it is an activator (*Fig. 7*). In *V. vulnificus*, H-NS was shown to repress *leuO* by binding to sites that overlap with ToxR and LeuO binding sites (67, 68). Our data also showed that *leuO* is an auto-repressor and that H-NS represses *leuO* in *V. parahaemolyticus*, which is likely using a similar regulatory mechanism to that in *V. vulnificus* (*Fig. 7*). Our growth data showed that deletion of either *leuO* or *hns* resulted in defects when grown at high salinity and a double Δ*leuO*/Δ*hns* mutant showed a severe growth defect that was more pronounced than either of the single mutants. The observed defects in high salinity suggests a larger role for both regulators in the osmotic stress response. The more pronounced growth defect observed for Δ*leuO*/Δ*hns* reinforces the hypothesis that these proteins interact at other loci important for increased salinity toleranace.

## Methods

### Bacterial strains, media and culture conditions

All strains and plasmids used in this study are listed in Table S2. *V. parahaemolyticus* RIMD2210633, a streptomycin-resistant clinical isolate, was used in this study as the wild-type (WT) strain (69, 70). Unless stated otherwise, *V. parahaemolyticus* was grown in either lysogeny broth (LB; Fisher Scientific, Fair Lawn, NJ) with 3% (wt./vole) NaCl (LB 3%) or M9 minimal media (47.8 mM Na_2_HPO_4_, 22 mM KH_2_PO_4_, 18.7 mM NH_4_Cl, 8.6 mM NaCl; Sigma Aldrich) supplemented with 2 mM MgSO_4_, 0.1 mM CaCl_2_, 20 mM glucose as the sole carbon source (M9G) and 3% (wt/vol) NaCl at 37°C. *E. coli* strains were grown in either LB supplemented with 1% (wt/vol) NaCl (LB1%) or M9G supplemented with 1% (wt/vol) NaCl. The strain *E. coli* β2155 *λpir* strain is a diaminopimelic acid (DAP) auxotrophic strain and was grown with 0.3 mM DAP (71). All strains were grown at 37°C with aeration. Antibiotics were added to growth media at the following concentrations as necessary: ampicillin (Amp), 100 µg/mL; chloramphenicol (Cm), 12.5 µg/mL; tetracycline (Tet), 1 µg/mL; kanamycin (Km), 50 µg/mL.

### Preparation of cellular extracts and proton nuclear magnetic resonance (^1^H-NMR)

Wild-type *V. parahaemolyticus* was grown overnight at 37°C in either M9G 1% NaCl or M9G 3%NaCl. Stationary phase cells were then pelleted and washed twice with 1XPBS. Three freeze-thaw cycles were performed with the cell pellets to increase lysis, and the cells were then suspended in 750 µL ethanol. Debris was pelleted by centrifugation, and the ethanol solution was transferred to a clean tube and evaporated under vacuum. Then deuterium oxide (D_2_O) was used to resuspend the pellet, and insoluble material was removed by centrifugation. The solution was then transferred to a 5-mm NMR tube for analysis on a Bruker AVANCE 600NMR spectrometer at a proton frequency of 600.13 MHz with a sweep of 12,376 Hz and a relaxation delay of 5 s. Sixteen scans were co-added for each spectrum.

### DNA-Affinity Pulldown

DNA-affinity chromatography pulldown was performed as previously described (72–74). The bait DNA primers were designed to amplify the 323-bp regulatory region of *ectA* (VP1722) with a biotin moiety added to the 5’ end. A 327-bp negative-control bait DNA probe was amplified from a coding region of *ectB* (VP1721), which was also labeled with a biotin moiety at the 5’ end. Both probes were amplified using Phusion HF polymerase (New England Biolabs) PCR. Ten reactions were completed for each probe and then pooled and purified using an ethanol extraction technique (75). The pulldown was completed twice, once after growth in non-inducing conditions (LB3%) and once after growth in inducing conditions (M9G3%) in an effort to obtain both positive and negative regulators of the ectoine biosynthesis operon. The wild-type strain was grown overnight in LB3%, cells were pelleted and washed two times with 1XPBS and diluted 1:50 into 500 mL of LB3% or M9G3%. The cultures were then grown to an OD_595_ of 0.5, pelleted and stored overnight at −80°C. The pellet was then suspended in 1.5 mL Fast-Break lysis buffer (Promega, Madison, WI) with 1 mM PMSF, 0.5 mM benzamidine, 0.1 µg/mL lysozyme and incubated for 30 min. at room temperature. The cells were then sonicated on ice at 30% pulse amplitude for 15 seconds with 1 min. rest, and this was repeated 5 times. The cell lysate was then clarified by centrifugation for 30 min. at 4°C. The cell lysate was precleared with streptavidin DynaBeads (Thermo Scientific, Waltham, MA) to remove nonspecific protein-bead interactions.

Streptavidin M280 DynaBeads (ThermoFisher) were then incubated two times with 10 µg of biotinylated probe for 20 min. The precleared cell lysate was then incubated for 30 min. with the bait-coated beads at room temperature with constant rotation in the presence of 100 µg sheared salmon sperm DNA as a non-specific competitor. This was completed twice with washes in between. Protein candidates were then eluted from the bait DNA-bead complex with a stepwise NaCl gradient (100 mM, 200 mM, 300 mM, 500 mM, 170 mM, and 1 M). Next, 6X SDS and 1 mM β-mercaptoethanol (BME) was added to elution samples which were then boiled at 95°C for 5 min. Samples were then separated using a 12% SDS-PAGE and visualized using the Pierce Silver Stain Kit (ThermoFisher). The P*_ectA_* bait and *ectB* negative control were loaded next to each other in order of increasing NaCl concentrations. Bands that were present in the P*_ectA_* bait but not the negative control lanes were excised, de-stained, and digested with trypsin following the standard procedure for mass spectrophotometry preparation with C18 ZipTips (Fisher Scientific, Fair Lawn, NJ). The samples were then individually analyzed with a Q-exactive Orbitrap mass spectrometer with nano-flow electrospray (Thermo). Proteins were subsequently identified using Proteome Discover 2.1 software.

### Protein Purification

Protein were purified as previously described (76). The full-length genes for *leuO* (VP0350) and *nhaR* (VP0527) were cloned into a pET-28a(+) vector with a C-terminal histidine tag, transformed into *E. coli* Dh5α, purified, and sequenced. These vectors pET*leuO* and pET*nhaR* were then transformed into *E. coli* BL21 (DE3) cells, and expression was induced with 0.5mM IPTG at 0.4 OD_595_ and then cells were grown overnight at 25°C. Cells were then pelleted and resuspended with lysis buffer (50 mM sodium phosphate, 200 mM NaCl, 20 mM imidazole, pH 7.4, 1.0 mM phenylmethanesulfonyl fluoride, 0.5 mM benzamidine) and lysed via sonication. Debris was pelleted, and supernatant applied to an IMAC column packed with HisPur Ni-NTA resin (ThermoFisher) that was equilibrated with column buffer (50 mM sodium phosphate, 200 mM NaCl, 20 mM imidazole, pH 7.4). The column was washed with buffer containing increasing concentrations of imidazole (20mM – 100mM) to remove any contaminants. His-tagged proteins were eluted using 500mM imidazole and then dialyzed overnight at 4°C in sodium phosphate buffer to remove any excess salts. Samples of supernatant and each flow through, wash, and elution were analyzed via SDS-PAGE gel to assess protein purity and verify purification.

### Electrophoretic mobility shift assay

DNA fragments were designed as outlined in Table S3, to encompass the entire region (322-bp) or sections of the ectoine biosynthesis regulatory region (125, 137, 106-bp). The concentration of purified protein (LeuO-His or NhaR-His) was determined using the Bradford Assay, and LeuO (0 to 2.175 uM) or NhaR (0 to 2.175 uM) was incubated for 20 minutes with 30 ng of each DNA fragment in a binding buffer (10 mM Tris [pH 7.4 at 4°C], 150 mM KCl, 0.1 mM dithiothreitol, 0.1 mM EDTA [pH 8.0], 5% polyethylene glycol). The reactions were then loaded (10 µL) on a 6% native acrylamide gel (pre-run 200V for 2 hours at 4°C) and run for 2 hours at 4°C (200V) in 0.5X TBE running buffer. To visualize, gels were stained in an ethidium bromide bath for 15 minutes.

### Mutant strain construction

In-frame deletion mutants of *nhaR* (VP0527), *torR* (VP1032), and *hns* (VP1133), were designed as previously described via allelic exchange (17). A truncated region of the *nhaR* (15-bp of the 891-bp), *torR* (54-bp of the 714-bp), and *hns* (33-bp of the 408-bp) genes was generated using the primers in Table S3. The truncated products were then ligated with the suicide vector pDS132 using the Gibson assembly protocol and transformed into *E. coli* Dh5α (77, 78). The resulting plasmids pDSΔ*nhaR*, pDSΔ*torR,* and pDSΔ*hns* were purified and transformed into *E. coli* β2155 λ*pir* (diaminopimelic acid auxotroph), followed by conjugation and homologous recombination into the *V. parahaemolyticus* RIMD2210633 genome. Single crossover of the plasmids into the genome were selected for by plating onto chloramphenicol (CM) and screened via PCR for a truncated allele. To induce a double-crossover event, the single-cross strain was grown overnight in the absence of CM, leaving behind either the truncated allele or the wild-type allele. The cultures were spread on 10% sucrose plates, and healthy colonies were screened for double-crossover and gene truncation; colonies that contain the plasmid will appear soupy on the plate due to the presence of the *sacB* selectable marker. In-frame deletions were confirmed with sequencing.

An in-frame deletion of *ompR* (VP0154) was created in *V. parahaemolyticus* RIMD2210633 through splicing by overlap extension (SOE) PCR and homologous recombination (79). Primers were designed to create a 692-bp truncated allele of *ompR* using the primer pair listed in Table S3. This truncated product was ligated into the cloning vector pJET1.2 with T4 DNA ligase and transformed into *E. coli* Dh5α λ*pir*. The truncated *ompR* allele was then excised from the pJETΔ*ompR* vector using restriction enzymes (SacI, XbaI) and ligated into the suicide vector pDS132. The pDSΔ*ompR* vector was then transformed into *E. coli* β2155 λ*pir* (DAP auxotroph), and the same procedure as described above was completed. Double mutants Δ*leuO/*Δ*hns* and Δ*hns/*Δ*nhaR* were generated by conjugating *E. coli* β2155 λ*pir* + pDS*hns* with *V. parahaemolyticus* Δ*leuO* and β2155 λ*pir* + pDS*nhaR* with *V. parahaemolyticus* Δ*hns.* The same procedure as described above was carried out to achieve these double deletions. To confirm that the phenotypes observed in the Δ*leuO,* Δ*hns* and Δ*nhaR* mutants were not due to secondary mutations within the genome, the strains were complemented with functional copies of the gene. The coding regions were amplified from *V. parahaemolyticus* RIMD2210633 genome and cloned into the expression vector pBAD33. Then transformation into *E. coli* β2155 was completed for both vectors. The expression vectors, pBAD*leuO,* pBAD*hns*, pBAD*nhaR* were then conjugated into the Δ*leuO,* Δ*hns* and Δ*nhaR* strains.

### Transcriptional GFP-reporter assay

The reporter construct pRUP*_ectA_-gfp* was created previously using the pRU1064 vector, which contains a promoterless *gfp* cassette, as well as Tet and Amp resistance genes (17, 80). Using *E. coli* β2155 λ*pir* containing pRUP*_ectA_-gfp*, the reporter plasmid was conjugated into each of the *V. parahaemolyticus* mutant strains Δ*leuO,* Δ*nhaR,* Δ*torR,* Δ*ompR,* Δ*hns,* Δ*leuO/*Δ*hns,* and Δ*hns/*Δ*nhaR.* Strains were grown overnight in LB3% with tetracycline. The cultures were then washed two times with 1XPBS, diluted 1:100 into M9G3% with tetracycline, and grown to OD ∼ 0.4 or OD ∼ 1.0 as indicated. The reporter expression was determined by measuring the relative fluorescence with excitation at 385 nm and emission at 509 nm in black, clear bottomed 96-well microplates on a Tecan Spark microplate reader with Magellan software (Tecan Systems, Inc., San Jose, CA). Specific fluorescence was calculated by dividing the relative fluorescence units (RFU) by the optical density of each well. At least two biological replicates were performed for each experiment. Statistics were completed either with a Student’s T-test or ANOVA followed by Tukey-Kramer Post Hoc test, as designated.

## ACKNOWLEDGEMENTS

This research was supported by a National Science Foundation grant (award IOS-1656688) to E.F.B. KBL and GJG were funded in part by a University of Delaware graduate fellowship award and KBL in part by a Chemistry-Biology Interface predoctoral training program grant: 5T32GM008550.

