## Supplemental Tables S1 to S3, Fig. S1 to S6 for "NhaR, LeuO and H-NS are part of an expanded regulatory network for ectoine biosynthesis expression"

**Table S1. Candidate proteins identified in DNA protein pulldown assay.** The mass spectrometry score provided is the combined sum of all scores for an observed spectra that can be matched to an individual peptide.

| Accession no. | Locus tag | Function/Predicted Domains | Score | Coverage | # AAs | MW [kDa] |
|---|---|---|---|---|---|---|
| Q87RV7 | VP0669 | Succinylglutamate/aspartoacyclase | 29.64 | 50.84 | 356 | 39.0 |
| Q87K62 | VPA0036 | N-terminal winged helix | 23.07 | 65.63 | 288 | 33.2 |
| Q87SD5 | VP0489 | Aerobic respiration control protein FexA | 17.15 | 9.66 | 238 | 26.9 |
| Q87S97 | VP0527 | NhaR Na+/H+ (NhaA) antiporter regulator | 15.06 | 38.18 | 296 | 33.9 |
| Q87SS2 | VP0350 | LeuO LysR family | 13.64 | 17.87 | 319 | 36.2 |
| Q87HC6 | VPA1039 | IcmF-related protein, type IV SS | 11.73 | 21.93 | 1172 | 133.7 |
| Q87QL6 | VP1133 | H-NS Nucleoid associated protein | 10.40 | 79.26 | 135 | 15.1 |
| Q87JF0 | VPA0303 | DeoR/GlpR family regulators | 9.94 | 17.89 | 246 | 26.9 |
| Q87M65 | VP2393 | GalR LacI family regulators | 7.87 | 3.31 | 332 | 36.2 |
| Q87QW7 | VP1032 | TorR DNA-binding response regulator | 7.74 | 15.19 | 237 | 26.7 |
| Q87FD8 | VPA1741 | MurR/RpiR family regulator | 7.47 | 4.95 | 283 | 30.8 |
| Q87IK0 | VPA0606 | Putative AraC-type regulator | 6.64 | 11.61 | 267 | 29.9 |
| Q87T18 | VP0252 | CytR LacI family regulator | 4.51 | 9.85 | 335 | 36.8 |
| Q87RT5 | VP0692 | GcvA LysR family regulator | 4.12 | 37.25 | 306 | 34.5 |
| Q87MQ9 | VP2172 | YheO, PAS and HTH domains | 3.92 | 40.87 | 252 | 28.2 |
| Q87N99 | VP1976 | MetR LysR family | 3.44 | 28.38 | 303 | 34.1 |
| Q87H16 | VPA1149 | ABC-transporter, SBP domain | 2.41 | 2.42 | 414 | 45.3 |
| Q87JP6 | VPA0202 | GGDEF family protein | 2.36 | 3.26 | 337 | 38.2 |
| Q87QW9 | VP1030 | PurR purine biosynthesis repressor | 2.35 | 4.19 | 334 | 37.7 |
| Q87TB6 | VP0154 | OmpR Osmolarity response regulator | 2.33 | 3.77 | 239 | 27.3 |
| Q87QB4 | VP1236 | Phosphogluconate repressor HexR | 2.33 | 3.52 | 284 | 31.5 |
| Q87FZ8 | VPA1522 | CrgA LysR family regulator | 2.30 | 3.56 | 309 | 34.4 |
| Q87S55 | VP0569 | PhoB DNA-bindng response regulator | 2.20 | 4.80 | 229 | 26.1 |
| Q87M71 | VP2387 | DeoR family transcriptional regulator | 2.15 | 3.80 | 263 | 29.2 |
| Q87J66 | VPA0387 | LysR family transcriptional regulator | 2.09 | 3.86 | 311 | 34.7 |
| Q87P35 | VP1683 | Hypothetical protein; periplasmic binding | 2.01 | 4.31 | 325 | 35.0 |

**Table S2. Strains and plasmids used in this study**

| Strain or plasmid | Genotype or description | Reference(s) or source |
|---|---|---|
| ***Vibrio parahaemolyticus*** | | |
| RIMD2210633 | O3:K6 clinical isolate, $Str^r$ | (Makino et al., 2003); |
| Δ*leuO* | RIMD2210633 ΔleuO (VP0350), $Str^r$ | (Whitaker et al., 2012) |
| Δ*nhaR* | RIMD2210633 ΔnhaR (VP0527), $Str^r$ | This study |
| Δ*hns* | RIMD2210633 Δhns (VP1133), $Str^r$ | This study |
| Δ*torR* | RIMD2210633 ΔtorR (VP1032), $Str^r$ | This study |
| Δ*ompR* | RIMD2210633 ΔompR (VP0154), $Str^r$ | This study |
| Δ*leuO*/Δ*hns* | RIMD2210633 ΔleuO Δhns (VP0350/VP1133), $Str^r$ | This study |
| Δ*hns*/Δ*nhaR* | RIMD2210633 Δhns ΔnhaR (VP0527/VP1133), $Str^r$ | This study |
| ***Escherichia coli*** | | |
| DH5α *λpir* | | ThermoFisher Scientific |
| B2155 *λpir* | Δ*dapA*::erm pir for bacterial conjugation | (Dehio & Meyer, 1997) |
| BL21 (DE3) | Protein expression strain | ThermoFisher Scientific |
| **Plasmids** | | |
| pDS132 | Suicide plasmid; CmR; *sacB*, R6Kg ori | (Philippe et al., 2004) |
| pDS*nhaR* | pDS132 harboring truncated *nhaR* allele | This study |
| pDS*torR* | pDS132 harboring truncated *torR* allele | This study |
| pDS*ompR* | pDS132 harboring truncated *ompR* allele | This study |
| pDS*hns* | pDS132 harboring truncated *hns* allele | This study |
| pRU1064 | promoterless-gfp UV, Ampr, Tetr; IncP origin | (Karunakaran et al., 2005) |
| pRUP*ectA* | pRU1064 with P*ectA*-gfp, Ampr, TetR | (Gregory et al., 2019) |
| pET28a (+) | Expression vector, 6xHis Tag, $Kan^r$ | Novagen |
| pET*leuO* | pET28a(+) harboring, *leuO*, $Kan^r$ | This study |
| pET*nhaR* | pET28a(+) harboring, *nhaR*, $Kan^r$ | This study |
| pBAD33 | Expression vector; *araB* promoter; CmR | (Bassler et al., 1997) |
| pBA*leuO* | pBAD33 with *leuO*, CmR | (Whitaker et al., 2012) |
| pBA*nhaR* | pBAD33 with *nhaR*, CmR | This study |
| pBA*hns* | pBAD33 with *hns*, CmR | This study |

**Table S3. Primers used in this study**

| Primer Name | Sequence (5′ – 3′) | Length (bp) |
|---|---|---|
| **Pull Down Probe Primers** | | |
| P*ectA*_PD fwd | Biotin-GCCACGACGACAAAACTAAC | 323 |
| P*ectA*_PD rev | CCAAGGTGCTGATGTGATCA | |
| *ectB*_PD fwd | Biotin-CGCTGTTCGAAACTATGCTG | 327 |
| *ectB*_PD rev | CGCCTGGTTTCCATTGATCA | |
| **Mutant Primers** | | |
| *nhaR* A gibson | accgcatgcgatatcgagctACAATACTTTTGCTTGGTCG | 553 |
| *nhaR* B gibson | tttactcaaaATGTGACATCGGCCAATTC | |
| *nhaR* C gibson | gatgtcacatTTTGAGTAAACCCGTATCGTTG | 490 |
| *nhaR* D gibson | gtggaattcccgggagagctGTCACGACGAAGCCAAGC | |
| *nhaR* FLF | AGACCACCTGCAACTAAGCC | 2157 |
| *nhaR* FLR | GCATTTCAGTCTGTGCGATCA | |
| *torR* A gibson | accgcatgcgatatcgagctGTTGAGCATCATTGCGGTC | 506 |
| *torR* B gibson | cttcaccgtgGACTAATACGTGATAGCTCATTTAC | |
| *torR* C gibson | cgtattagtcCACGGTGAAGGCTACATG | 502 |
| *torR* D gibson | gtggaattcccgggagagctTGATGGTGGCTCTGAGATC | |
| *torR* FLF | CCTGAACCCCAAAATCTGCC | 2033 |
| *torR* FLR | ACCAAGCTTCTCATGCCACT | |
| *ompR* A gibson | TCTAGAGCGCCGAAAGTTAGATGAAG | 301 |
| *ompR* B gibson | CAGCTGAGATCTGGTACCCCTGCATTGAAACATCCTTT | |
| *ompR* C gibson | GGTACCAGATCTCAGCTGCCAGACGGCAAAGAGTCGTAA | 336 |
| *ompR* D gibson | GAGCTCTAATAACTGGCGACGCAATG | |
| *ompR* FLF | TGGATTAAAACGAGCCCTTG | 1329 |
| *ompR* FLR | TCGATGTGCATCCACAAAAT | |
| *hns* A gibson | accgcatgcgatatcgagctTGCGCCTCTACGGCAACAAC | 654 |
| *hns* B gibson | cgtctagtgaCTCTGACATAACGATTCCTATTAATAGGTTAAC | |
| *hns* C gibson | tatgtcagagTCACTAGACGATTTCCTAATCTAATC | 609 |
| *hns* D gibson | gtggaattcccgggagagctGGCGGGTTACACTGAAGAC | |
| *hns* FLF | GCAATCTTCGCCTTCCACTA | 1699 |
| *hns* FLR | AAGACGCTAGAGTGGGCTGA | |

**Continued Table S3. Primers used in this study**

| **Protein Expression Primers** | | |
|---|---|---|
| *leuO* fwd | tggacagcaaatgggtcgcgCAATGTTAGAGAAGAAAGATG | 1002 |
| *leuO* rev | gtcgacggagctcgaattcgTTATTTTGATGCGACCAC | |
| *nhaR* fwd | tggacagcaaatgggtcgcgCGATGTCACATCTAAATTACAAC | 933 |
| *nhaR* rev | gtcgacggagctcgaattcgTTACTCAAACATTTGACTGAAATC | |
| **EMSA Primers** | | |
| P*ectA*Fwd_2 | CCAAGGTGCTGATGTGATCA | 323 |
| P*ectA*Rev_2 | GTTAGTTTTGTCGTCGTGGC | |
| P*ectA*Fwd_1A | CCAAGGTGCTGATGTGATCA | 125 |
| P*ectA*Rev_1A | CACATTAATCCAGATTAAAACGCAG | |
| P*ectA*Fwd_1B | CTGCGTTTTAATCTGGATTAATGTG | 137 |
| P*ectA*Rev_1B | CCCACTGCATTCTGACTCAT | |
| P*ectA*Fwd_1C | ATGAGTCAGAATGCAGTGGG | 106 |
| P*ectA*Rev_1C | CCACGACGACAAAACTAAC | |
| P*leuO*Fwd | GTTTGTTTGCTCGGATTGTT | 597 |
| P*leuO*Rev | TGAGTGCGCCTCTTTTGTAA | |

### Ectoine Biosynthesis Operon

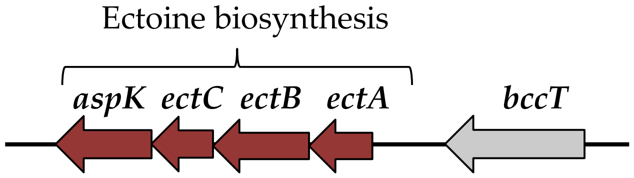


### Ectoine Biosynthesis Pathway

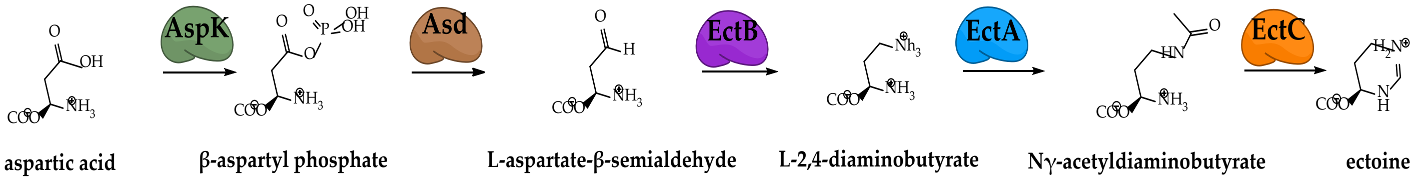


**Fig. S1. Ectoine biosynthesis. A.** Ectoine biosynthesis operon **B.** Biosynthesis of ectoine (1,4,5,6-tetrahydro-2-methyl-4-pyrimidinecarboxylic acid) by EctA, EctB, EctC, and aspartokinase (*aspk*) from aspartic acid.

**Supplemental Fig. S1**

**A. H-NMR analysis of ectoine biosynthesis**

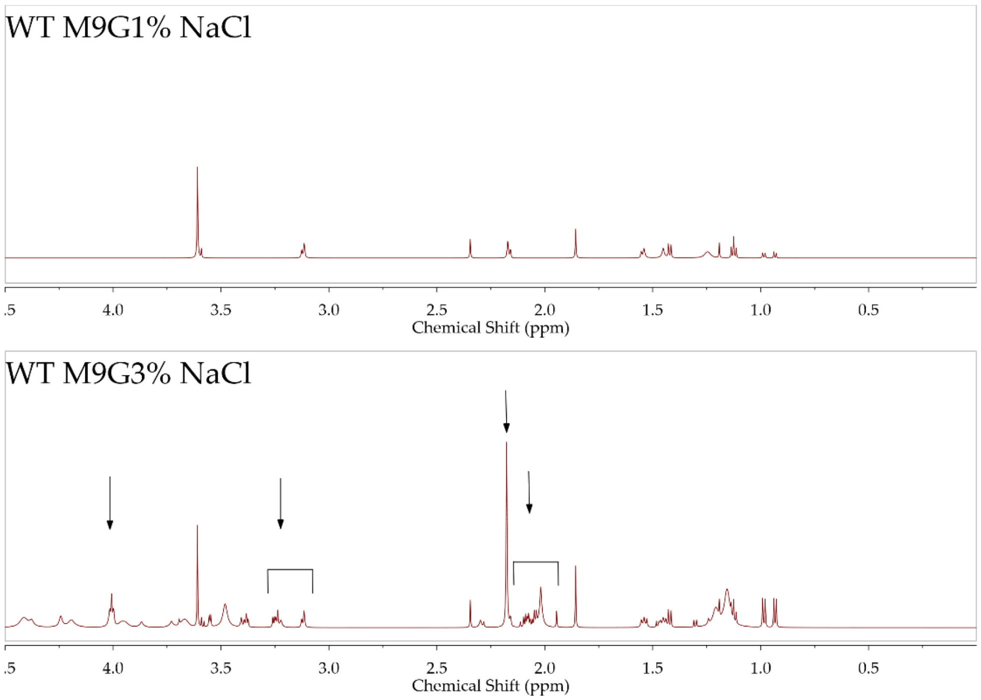


**B. P*ectA-gfp* expression increased in M9G 3% NaCl**

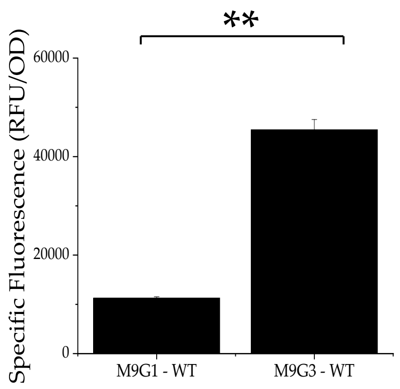


**Fig. S2. Ectoine production and P*ectA-gfp* expression induced in M9G3% NaCl A.**[1]H-NMR spectroscopy of *V. parahaemolyticus* wild type (WT) cell lysate grown in M9G 1% NaCl or M9G 3% NaCl. The spectral peaks for ectoine are indicated with arrows, and chemical shifts are expressed in ppm. **B**. Expression of P*ectA-gfp* transcriptional fusion in *V. parahaemolyticus* WT. Cultures were grown overnight in M9G 1% NaCl or M9G 3% NaCl and relative fluorescence intensity (RFU) was measured. Specific fluorescence was calculated by dividing RFU by OD. Mean and standard deviation of two biological replicates are shown. Statistics were calculated using a Student's t-test; (**, $P < 0.005$).

**Supplemental Fig. S2**

**A. DNA-Protein Pull Down – protein separation with SDS-PAGE**

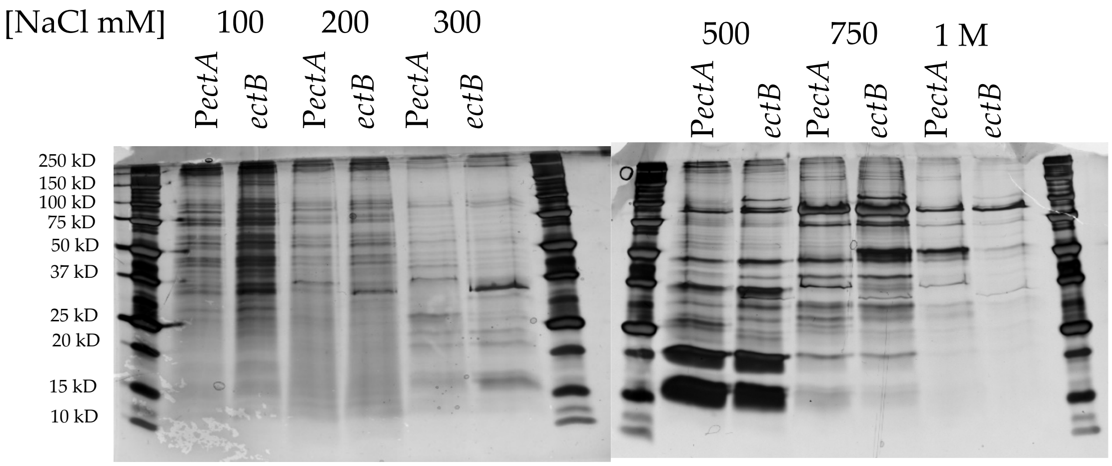


**B. Bands excised for analysis**

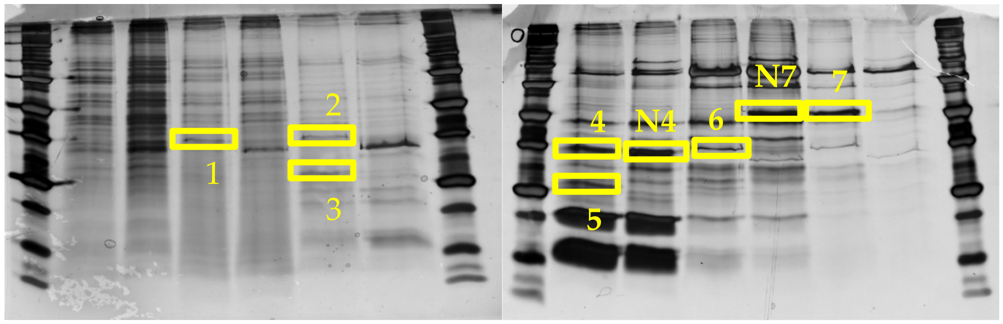


**Fig. S3. DNA-affinity pulldown, protein separation with SDS-PAGE. A**. Proteins that bind to the regulatory region of *ectABC-aspk* were identified with DNA affinity chromatography and mass spectrometry. **B**. Shown here are gel pieces that were excised and independently analyzed to identify candidate proteins labelled 1, 2, 3, 5, and 6. Negative controls are labelled N4 and N7.

**Supplemental Fig. S3**

**Protein Purification – LeuO & NhaR**

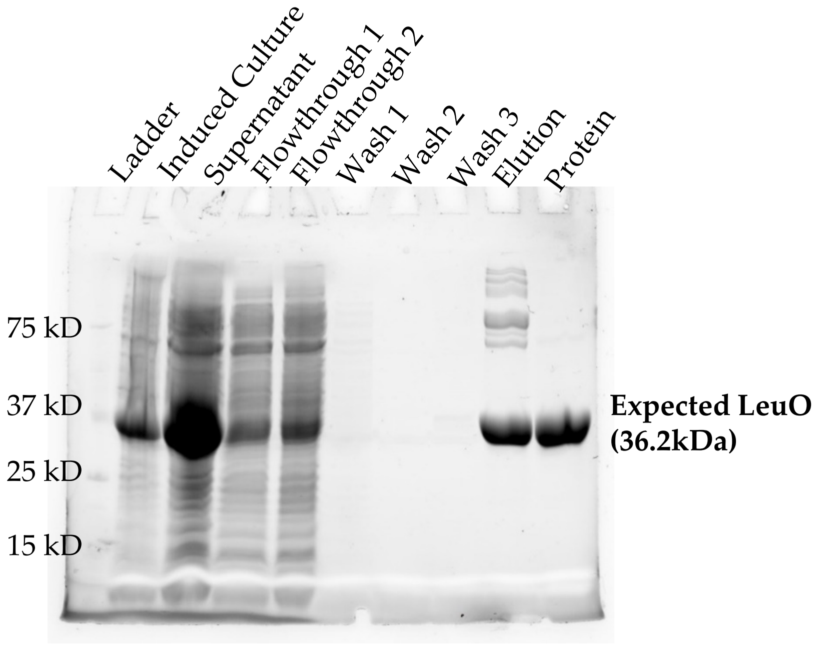


LeuO Protein Purification

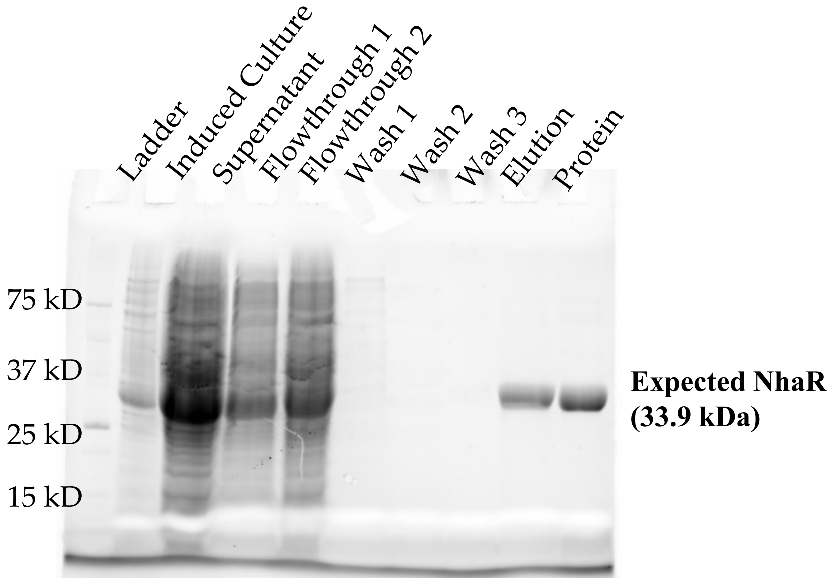


NhaR Protein Purification

**Fig. S4. LeuO and NhaR protein purification.** Proteins tagged with 6xHis purified, analyzed in a SDS-PAGE gel.

**Supplemental Fig. S4**

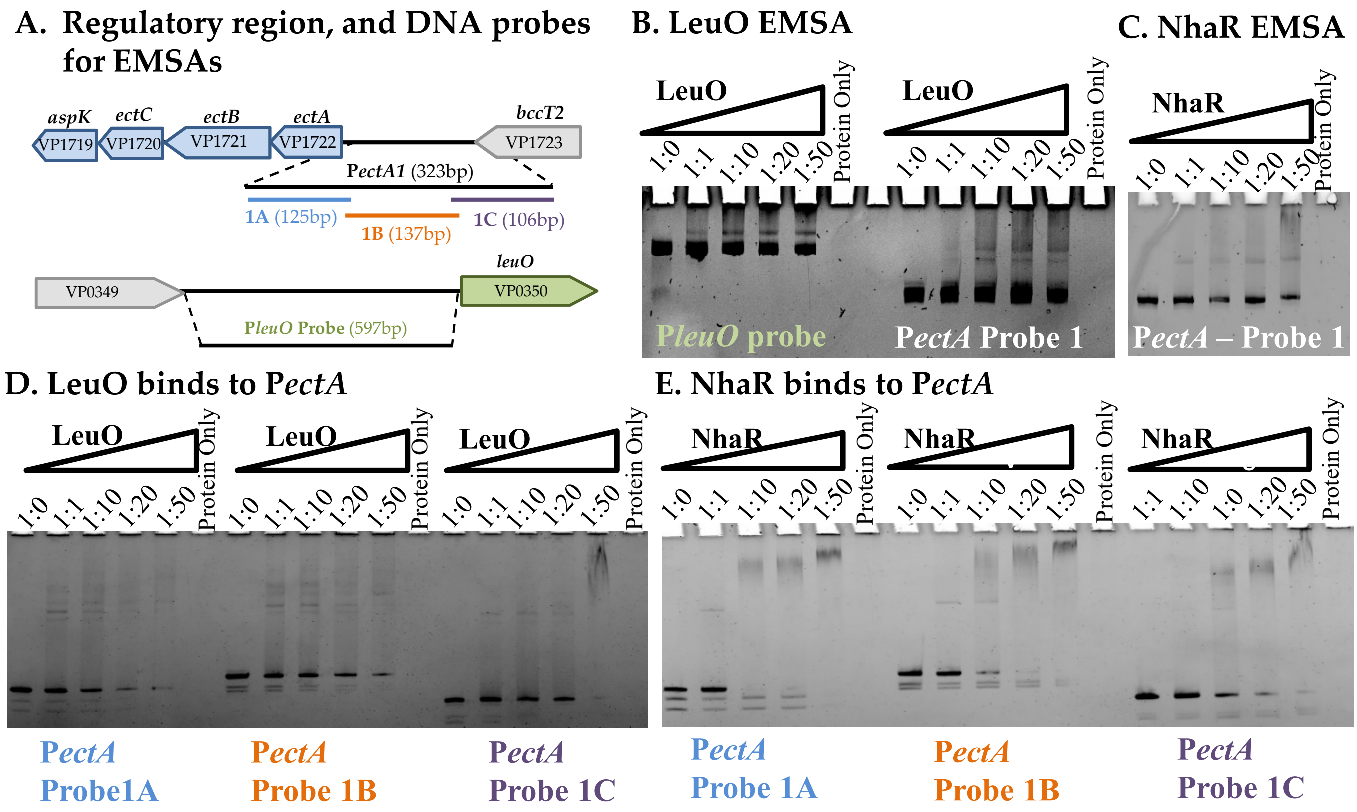


**Figure S5. LeuO and NhaR bind to the *ectABC-ask* regulatory region. A.** Region upstream of *ectABC-aspk* operon ($P_{ectA}$) is presented here both as a large probe (2) and segmented into three smaller probes (1A, 1B, and 1C) and upstream region of *leuO*. **B.** LeuO binds to probe $P_{ectA}$-2 and to the *leuO* regulatory region. **C.** NhaR binds to the $P_{ectA}$-2 regulatory region. **D**. LeuO binds to segmented portions of $P_{ectA}$ probes. **E.** NhaR binds to segmented portions of $P_{ectA}$ probes. EMSA's were performed with 30 ng of $P_{ectA}$-*gfp* probe and purified LeuO or NhaR protein (0 to 2.175 uM) with DNA: protein molar rations of 1:0, 1:1, 1:10, 1:20, and 1:50

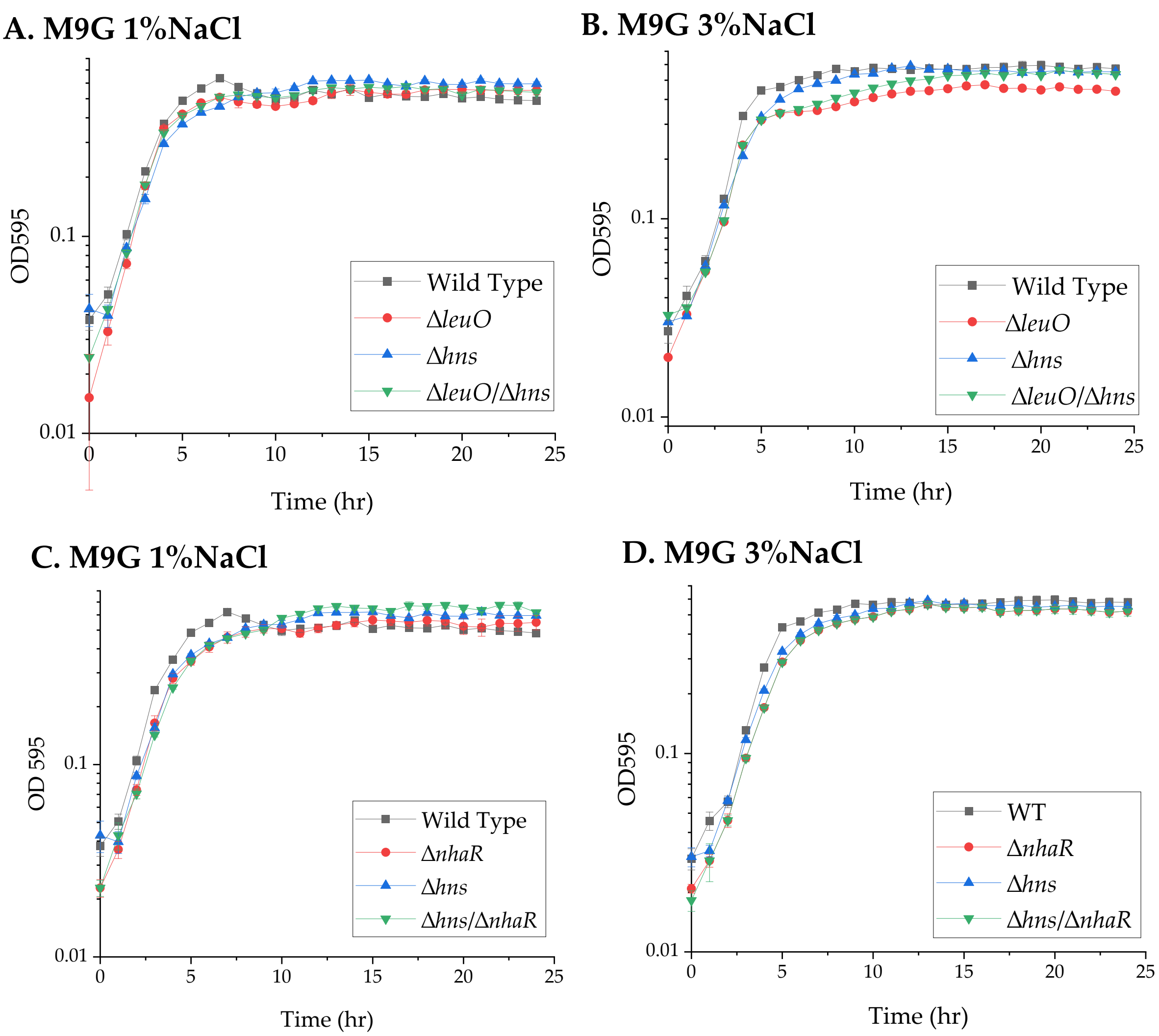


**Figure S6. Growth analysis of mutants in 1%NaCl and 3%NaCl. A.** Wild type, *ΔleuO, Δhns*, and *ΔleuO/Δhns* in M9G supplemented with 1%NaCl or **B.** M9G 3%NaCl. **C.** Wild type, *ΔnhaR, Δhns*, and *Δhns/ΔnhaR* in M9G 1%NaCl or **D.** M9G 3%NaCl. Cells were grown to OD 0.5 in M9G1% and then inoculated into M9G 1%NaCl or M9G 3%NaCl and growth was measured every hour for 24 h at 37°C. Mean and standard deviation of two biological replicates are shown.

**Supplementary Fig. S6**
